# Neurosecretory protein GL-induced fat accumulation is accompanied by repressing the immune-inflammatory response in the adipose tissue of mice

**DOI:** 10.1101/2021.08.27.457926

**Authors:** Keisuke Fukumura, Yuki Narimatsu, Eiko Iwakoshi-Ukena, Megumi Furumitsu, Hidemasa Bono, Kazuyoshi Ukena

**Affiliations:** Laboratory of Neurometabolism, Graduate School of Integrated Sciences for Life, Hiroshima University, Hiroshima, Japan; Laboratory of Genome Informatics, Graduate School of Integrated Sciences for Life, Hiroshima University, Hiroshima, Japan

## Abstract

We have recently identified neurosecretory protein GL (NPGL), a small secretory protein expressed in the vertebrate hypothalamus, as an orexigenic factor with remarkable fat accumulation by overexpression of the NPGL precursor gene (*Npgl*) for two months. In the present study, we analyzed the effects of short-term *Npgl* overexpression for 18 days as the early stage of obesity to address the mechanisms underlying obese-like phenotype. Similar to previous studies, short-term *Npgl* overexpression stimulated food intake and fat accumulation in the white adipose tissues (WAT), whereas the masses of the brown adipose tissue, testis, liver, heart, and muscle remained unchanged. In addition, we observed increased blood insulin and leptin levels due to *Npgl* overexpression, while little changes were induced in blood glucose, free fatty acids, triglyceride, and cholesterol levels. Furthermore, transcriptome analysis of the inguinal WAT using RNA-sequencing technique revealed that overexpression of *Npgl* upregulated the genes involved in cytoskeleton regulation, whereas it decreased those involved in immune-inflammatory responses. These results suggest that NPGL plays a crucial role in enlarging adipocytes and suppressing inflammation to avoid metabolic abnormalities, eventually contributing to accelerating energy storage.

## Introduction

Although obesity has become a public health problem worldwide, there are no definitive therapeutic approaches. Genetic and environmental factors are closely associated with obesity and its comorbidities, such as depression, type 2 diabetes, and cardiovascular disease^1–3^. As obesity progresses, excess fat accumulation induces chronic inflammation in adipose tissue, which is regulated by adipose tissue-resident immune cells, such as T cells, B cells, and macrophages^4–6^. Indeed, these cells in adipose tissue correlated with total adiposity and adipose cell size, and secreted the majority of inflammatory cytokines, causing metabolic abnormalities in obesity^7–10^. On the other hand, as it is known that continuous overfeeding can contribute to obesity development, the mechanisms controlling feeding behavior and systemic metabolism have been explored^11–13^. So far, hypothalamic neuropeptides involved in feeding behavior and metabolism have been identified, including potent orexigenic and anorexigenic factors such as neuropeptide Y (NPY), agouti-related peptide (AgRP), and proopiomelanocortin (POMC)- derived α-melanocyte-stimulating hormone^11–13^. In addition, peripheral factors involved in regulation of feeding behavior have been discovered. Ghrelin, an orexigenic peptide produced by the stomach, evokes feeding behavior via the hypothalamic NPY/AgRP circuit^14–16^. Conversely, leptin, an anorexigenic peptide, is secreted from the white adipose tissue (WAT) and affects NPY/AgRP and POMC neurons^17–19^.

Throughout our investigations of the regulatory mechanisms of energy homeostasis, we have previously identified a novel cDNA encoding a peptide hormone precursor in the chick hypothalamus^20^. Deduced precursor protein, which included a small secretory protein of 80 amino acids with a Gly-Leu-NH_2_ sequence at the C-terminus, has been named neurosecretory protein GL (NPGL)^20^. Homologous NPGL proteins have been discovered in mammals, including humans, rats, and mice, implying that NPGL possesses a vital physiological function across species^21^. Indeed, intracerebroventricular (i.c.v.) infusion of NPGL increases food intake and affects energy metabolism in avian species^22,23^. Likewise, we have also shown that overexpression of the NPGL precursor gene (*Npgl*) elicits food intake and subsequent fat accumulation in the WAT of rats through *de novo* lipogenesis using dietary carbohydrates^24^. Using mice, we have demonstrated that acute i.c.v. infusion of NPGL increases food intake and that chronic i.c.v. infusion of NPGL increases food intake and results in considerable fat accumulation in adipose tissue^25,26^. Furthermore, we have recently shown that *Npgl* overexpression for two months elicits obesity phenotypes, such as increased food intake and considerable fat accumulation in mice^27^. Our data has also revealed that *Npgl* overexpression hardly induces metabolic abnormalities, such as hyperglycemia and hyperlipidemia^27^. However, the underlying mechanisms of fat accumulation and metabolic normality in *Npgl*-overexpressing mice are poorly understood.

In this study, we performed short-term *Npgl* overexpression for 18 days as the early stage of obesity development to address the mechanisms underlying fat accumulation and avoid secondary effects caused by prolonged gene overexpression. In addition, we conducted transcriptome analysis of the inguinal WAT (iWAT) using RNA sequencing (RNA-seq) techniques to uncover the molecular basis in obesity phenotypes induced by NPGL.

## Results

### Effects of NPGL-precursor gene overexpression for 18 days on food intake and body mass

To investigate the effects of NPGL on energy homeostasis as the early stage of obesity development in mice, we performed NPGL-precursor gene (*Npgl*) overexpression for 18 days. The results showed that *Npgl* overexpression increased cumulative food intake and body mass within 18 days (Fig. 1A and B).

**Figure 1.**
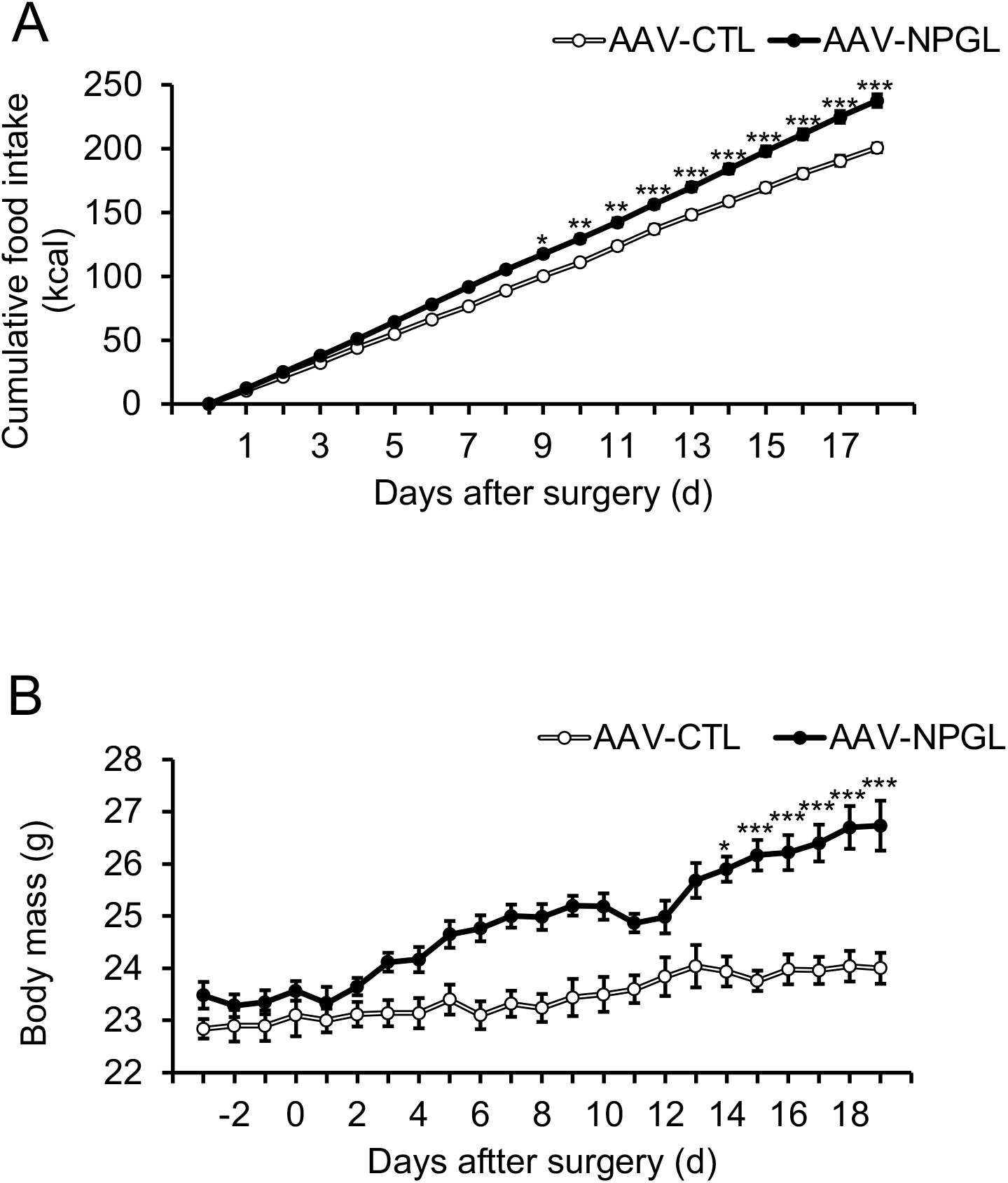
Effects of *Npgl* overexpression for 18 days on food intake and body mass. These mice were injected with an adeno-associated virus (AAV) vector, either a control (AAV-CTL) or a vector carrying the NPGL precursor gene (AAV-NPGL). (**A**) Cumulative food intake and (**B**) body mass. Each value represents the mean ± standard error of the mean (n = 5–6; \**p* < 0.05, \*\**p* < 0.01, \*\*\**p* < 0.005 by Student’s *t*-test).

### Effects of NPGL-precursor gene overexpression for 18 days on body composition and blood biomarkers

After observing increased food intake and body mass within 18 days, we next investigated the effects of *Npgl* overexpression on body composition. Measurement of tissue and organ masses revealed that *Npgl* overexpression increased the masses of the inguinal, epididymal, retroperitoneal, and perirenal WAT, whereas the interscapular brown adipose tissue mass was not changed (Fig. 2A and B). *Npgl* overexpression did not affect the mass of the gastrocnemius muscle, the muscle in the calf of the leg (Fig. 2C). Moreover, *Npgl* overexpression increased the mass of the kidney, despite no differences in the masses of the testis, liver, and heart (Fig. 2D).

**Figure 2.**
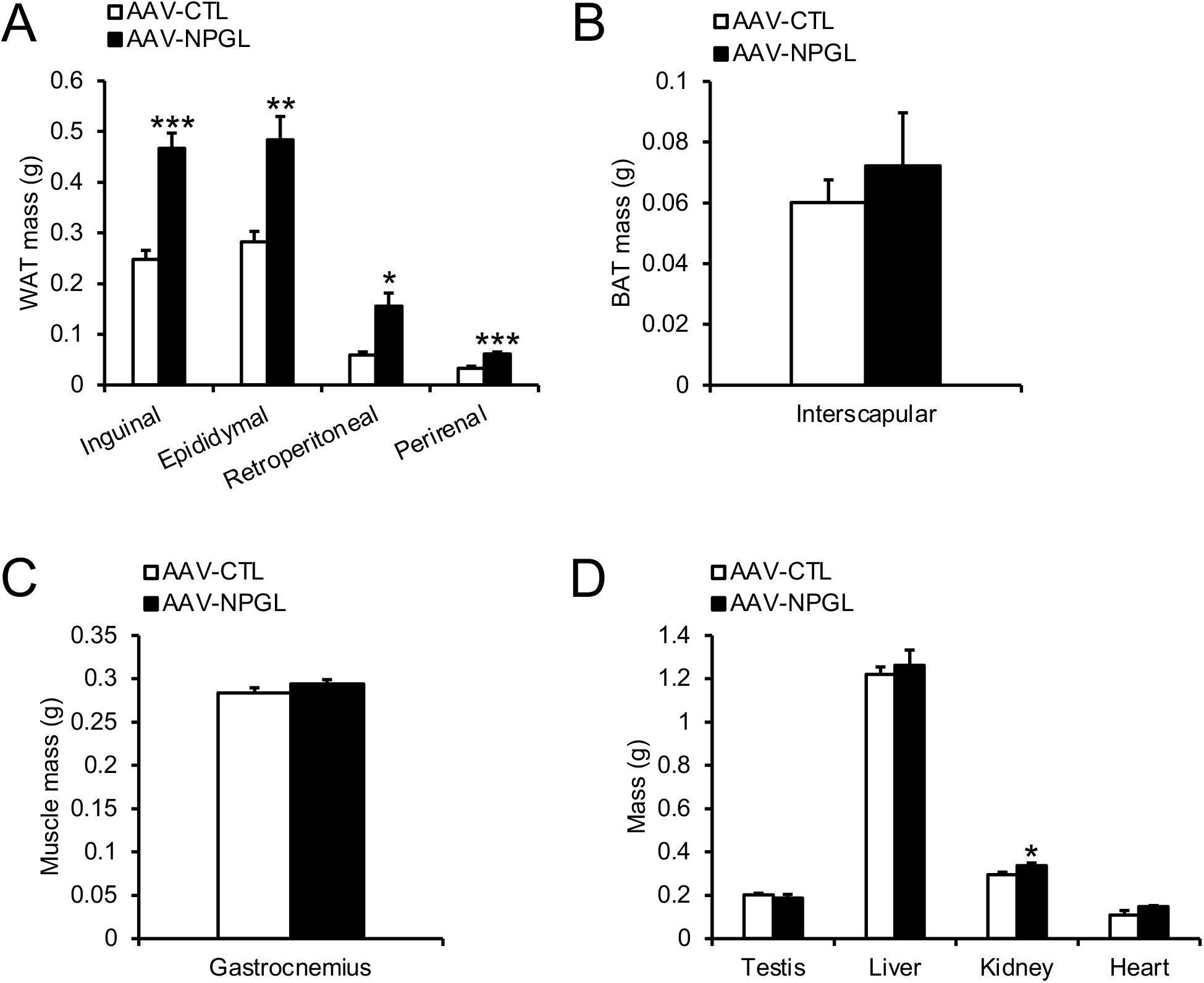
Effects of *Npgl* overexpression for 18 days on tissue and organ masses. These mice were injected with an adeno-associated virus (AAV) vector, either a control (AAV-CTL) or a vector carrying the NPGL precursor gene (AAV-NPGL). (**A**) Masses of the inguinal, epididymal, retroperitoneal, and perirenal white adipose tissue (WAT). (**B**) The interscapular brown adipose tissue (BAT) mass. (**C**) The gastrocnemius muscle mass. (**D**) Masses of the testis, liver, kidney, and heart. Each value represents the mean ± standard error of the mean (n = 5–6; \**p* < 0.05, \*\**p* < 0.01, \*\*\**p* < 0.005 by Student’s t-*t*est).

Regarding blood biomarkers, *Npgl* overexpression increased blood insulin and leptin levels but did not affect blood levels of glucose, free fatty acid, triglyceride, and cholesterol (Fig. 3).

**Figure 3.**
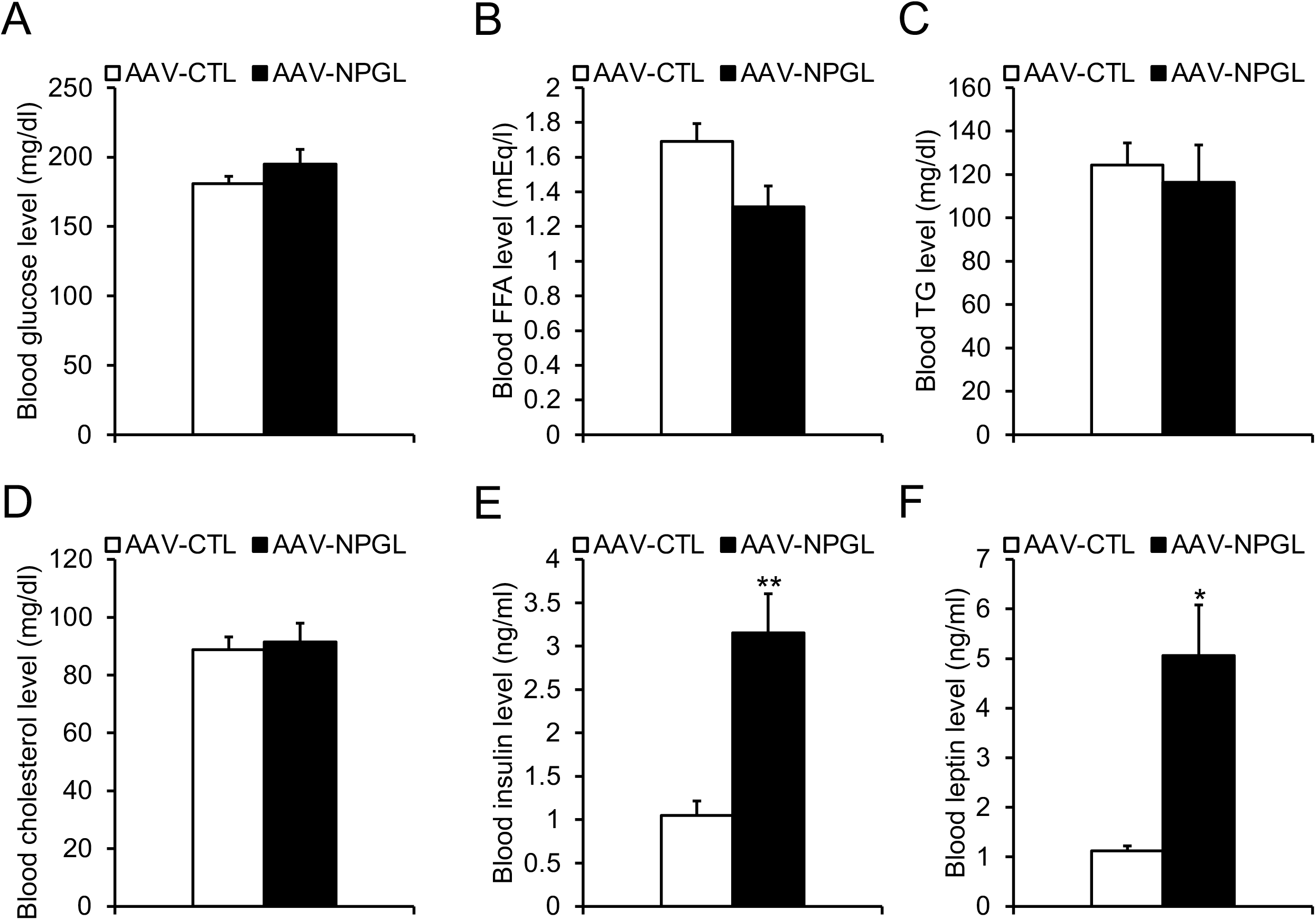
Effects of *Npgl* overexpression for 18 days on blood biomarkers. These mice were injected with an adeno-associated virus (AAV) vector, either a control (AAV-CTL) or a vector carrying the NPGL precursor gene (AAV-NPGL). (**A–F**) Levels of blood glucose (**A**), triglycerides (TG) (**B**), free fatty acids (FFA) (**C**), cholesterol (**D**), insulin (**E**), and leptin (**F**). Each value represents the mean ± standard error of the mean (n = 5–6; \**p* < 0.05, \*\**p* < 0.01 by Student’s *t*-test).

### Effects of NPGL-precursor gene overexpression for 18 days on mRNA expression of lipid metabolism-related genes and lipogenic activity

To address the mechanisms underlying increased fat accumulation in the WATs of *Npgl*-overexpressing mice, we subsequently measured mRNA expression of genes involved in lipid metabolisms in the iWAT and liver by quantitative RT-PCR (qRT-PCR). We analyzed the following: acetyl-CoA carboxylase (*Acc*), fatty acid synthase (*Fas*), stearoyl-CoA desaturase 1 (*Scd1*), and glycerol-3-phosphate acyltransferase 1 (*Gpat1*) as lipogenic enzymes; carbohydrate-responsive element-binding protein α (*Chrebpα*) and carbohydrate-responsive element-binding protein β (*Chrebpβ*) as lipogenic transcription factors; carnitine palmitoyl transferase 1a (*Cpt1a*), adipose triglyceride lipase (*Atgl*), and hormone-sensitive lipase (*Hsl*) as lipolytic enzymes; glyceraldehyde-3-phosphate dehydrogenase (*Gapdh*) as a carbohydrate metabolism enzyme; solute carrier family 2 member 4 (*Slc2a4*) as a glucose transporter; cluster of differentiation 36 (*Cd36*) as a fatty acid transporter. qRT-PCR showed that *Npgl* overexpression for 18 days increased mRNA expression levels of *Scd1, Gpat1, Chrebpα, Cpt1a, Atgl, Hsl, Gapdh, Slc2a4*, and *Cd36* in the iWAT (Fig. 4A). In the liver, mRNA expression levels of *Chrebpα* and *Gapdh* were upregulated in *Npgl*-overexpressing mice (Fig. 4B).

**Figure 4.**
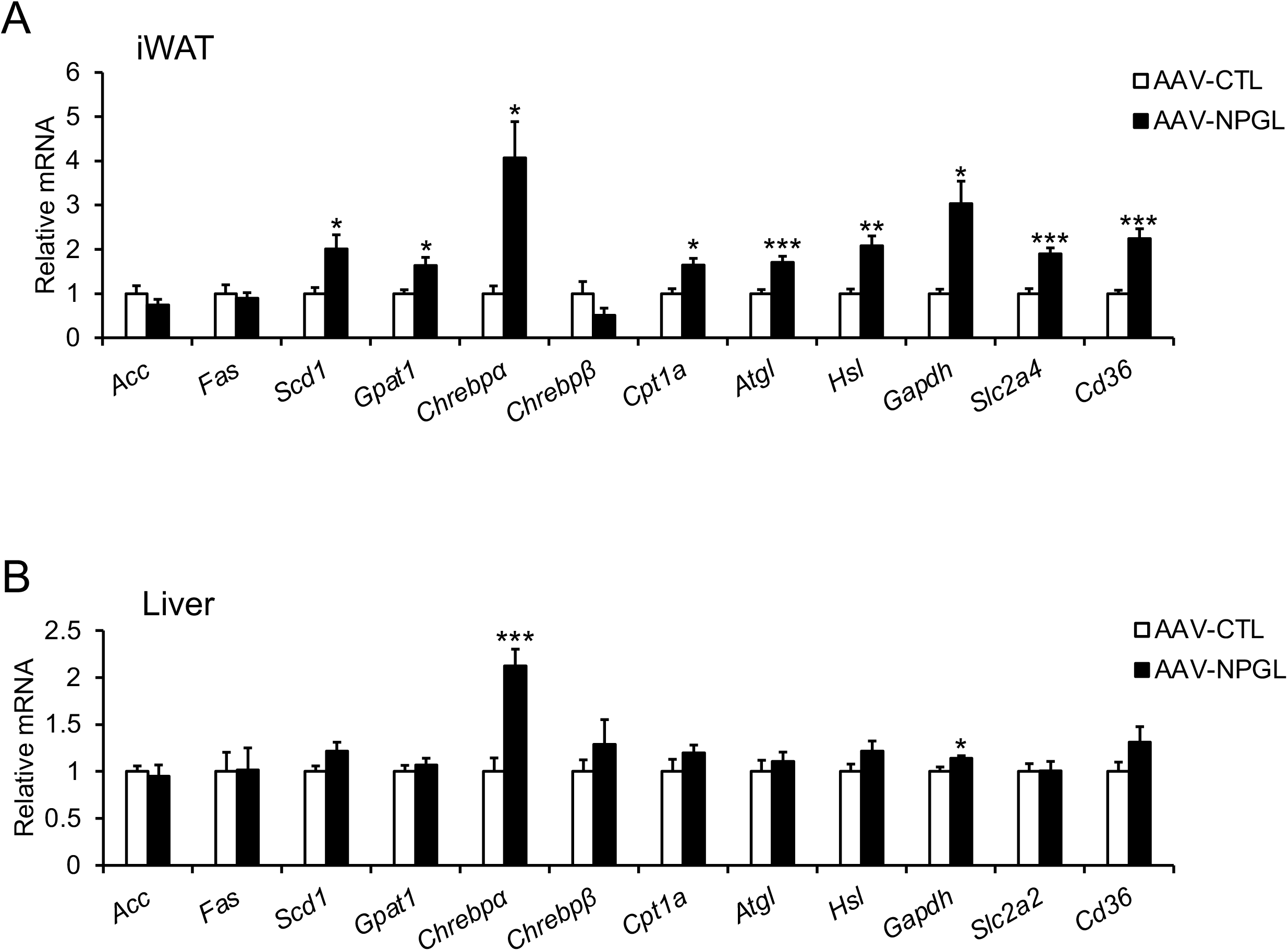
Effects of *Npgl* overexpression for 18 days on mRNA expression of genes related to lipid metabolism. These mice were injected with an adeno-associated virus (AAV) vector, either a control (AAV-CTL) or a vector carrying the NPGL precursor gene (AAV-NPGL). (**A, B**) mRNA expression levels in the inguinal white adipose tissue (iWAT) (**A**) and liver (**B**). Each value represents the mean ± standard error of the mean (n = 5–6; \**p* < 0.05, \*\**p* < 0.01, \*\*\**p* < 0.005 by Student’s *t*-test).

To analyze the activity of lipogenic factor in the iWAT, we measured the fatty acid ratio using gas chromatography-mass spectrometry (GC-MS). The ratios of palmitoleate to palmitate (16:1/16:0) and oleate to stearate (18:1/18:0) evaluate the enzymatic activity of stearoyl-CoA desaturase 1 (SCD1)^28^. The ratio of 16:0/18:2n-6 is an index of *de novo* lipogenesis^29,30^. GC-MS analysis showed that *Npgl* overexpression for 18 days had little effect on the ratios of 16:1/16:0, 18:1/18:0, and 16:0/18:2n-6 in the iWAT (Supplemental fig. 1). These data indicated that *Npgl* overexpression for 18 days does not promote *de novo* lipogenesis, even though it increased the mRNA expressions of the gene involved in lipid metabolism.

### Transcriptome analysis of the iWAT of *Npgl*-overexpressing mice

To explore the molecular basis of obese-like phenotypes in *Npgl*-overexpressing mice for 18 days, we next conducted transcriptome analysis using the RNA-seq technique. Using TCC-GUI^31^, 883 differential expressed genes (DEGs) between the iWAT of control and *Npgl*-overexpressing mice were screened based on the criteria of false discovery rate (FDR) < 0.1. MA plots showed a broad overview of changes in gene expression between control and *Npgl*-overexpressing mice (Fig. 5A). Of 883 DEGs, 682 genes were upregulated, while 201 genes were downregulated by *Npgl* overexpression (Fig. 5B). Gene ontology (GO) enrichment analysis was performed with upregulated and downregulated DEGs, respectively. The results showed that *Npgl* overexpression upregulated the genes involved in muscle structure development (GO:0061061), muscle system process (GO:0003012), and striated muscle cell development (GO:0055002), as well as others (Fig. 6A). Indeed, we found that *Npgl* overexpression increased mRNA expressions related to cytoskeleton regulation, such as plakophilin 2 (*Pkp2*) and Tublin alpha 1A (*Tuba1a*) (Fig. 6B). Furthermore, *Npgl* overexpressing remarkably upregulated mRNA expression of Mesoderm specific transcript (*Mest*), a gene related to adiposity.

**Figure 5.**
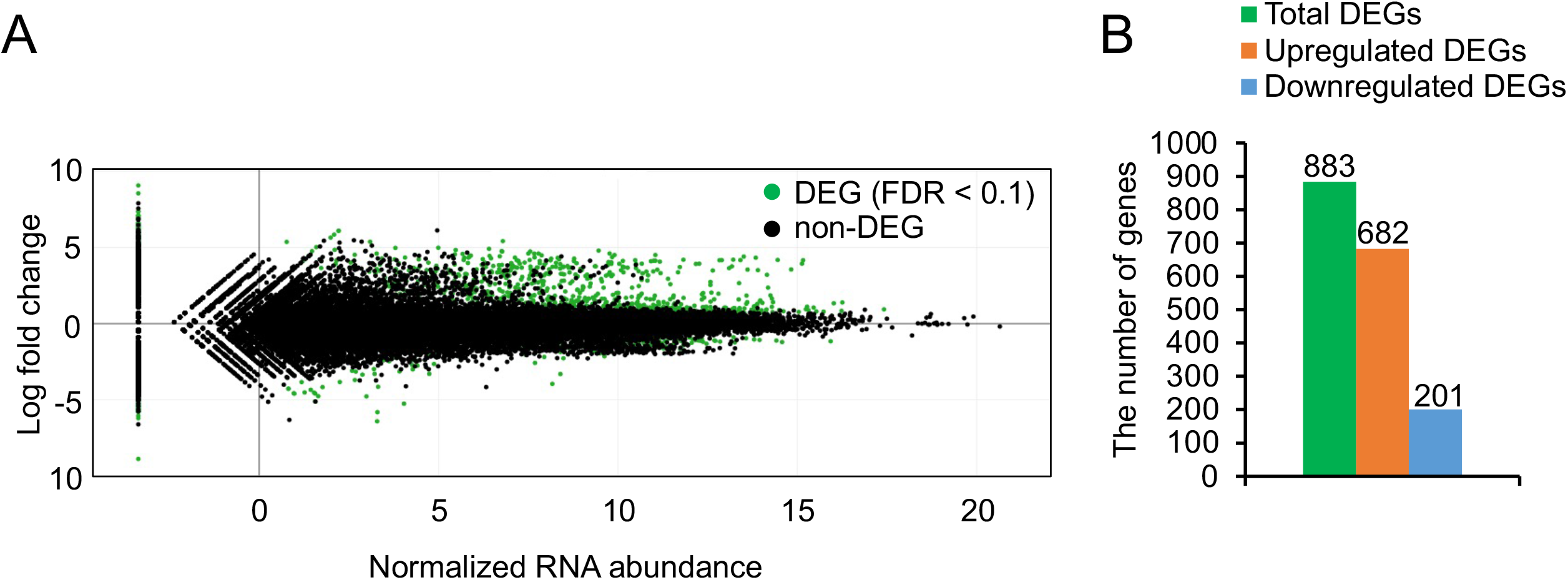
Differentially expressed genes (DEGs) in the inguinal white adipose tissue (iWAT) of *Npgl* overexpressing mice. (A) MA plots analysis showing DEGs between the iWAT of *Npgl* overexpressing mice and control mice. Green dots indicate DEGs with statistical significance based on false discovery rate (FDR) < 0.1. (B) The number of upregulated and downregulated DGEs identified.

**Figure 6.**
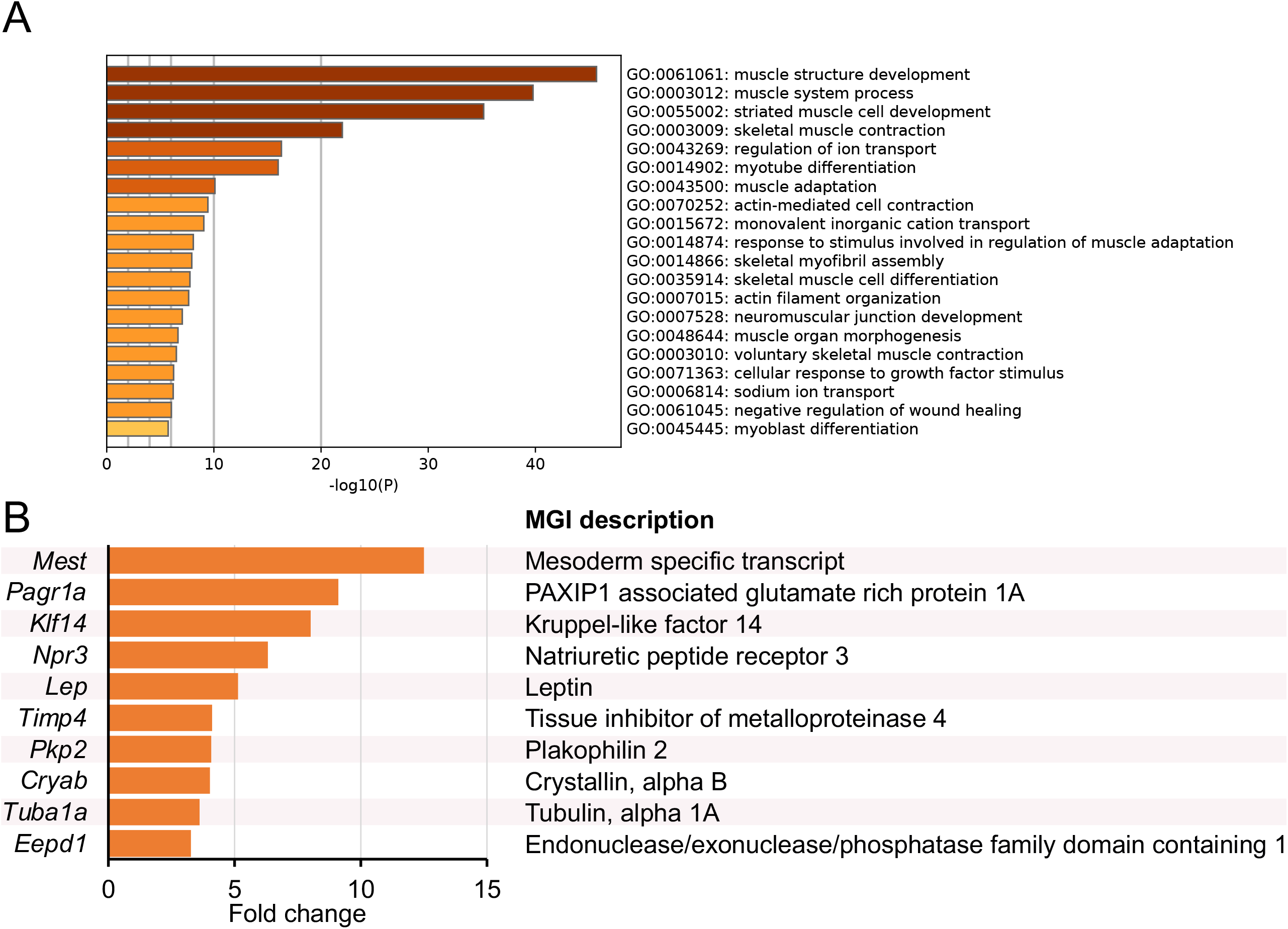
Gene ontology (GO) enrichment analysis of upregulated DEGs. (**A**) A heat map of GO terms across the upregulated DEGs list, colored to indicate the p values. Meta scape were used for this analysis. (**B**) Selected 10 genes indicating low false discovery rate of upregulated DEGs.

Concerning downregulated DEGs, GO enrichment analysis showed that *Npgl* overexpression decreased mRNA expressions of the genes involved in adaptive immune response (GO:0002250), defense response to other organism (GO:0098542), and regulation of cytokine production (GO: 0001817), as well as others (Fig. 7A). We found that *Npgl* overexpression remarkably downregulated mRNA expression of the genes related to inflammation and lymphocyte activation, such as tumor necrosis factor (ligand) super family member 18 (*Tnfsf18*) (Fig. 7B).

**Figure 7.**
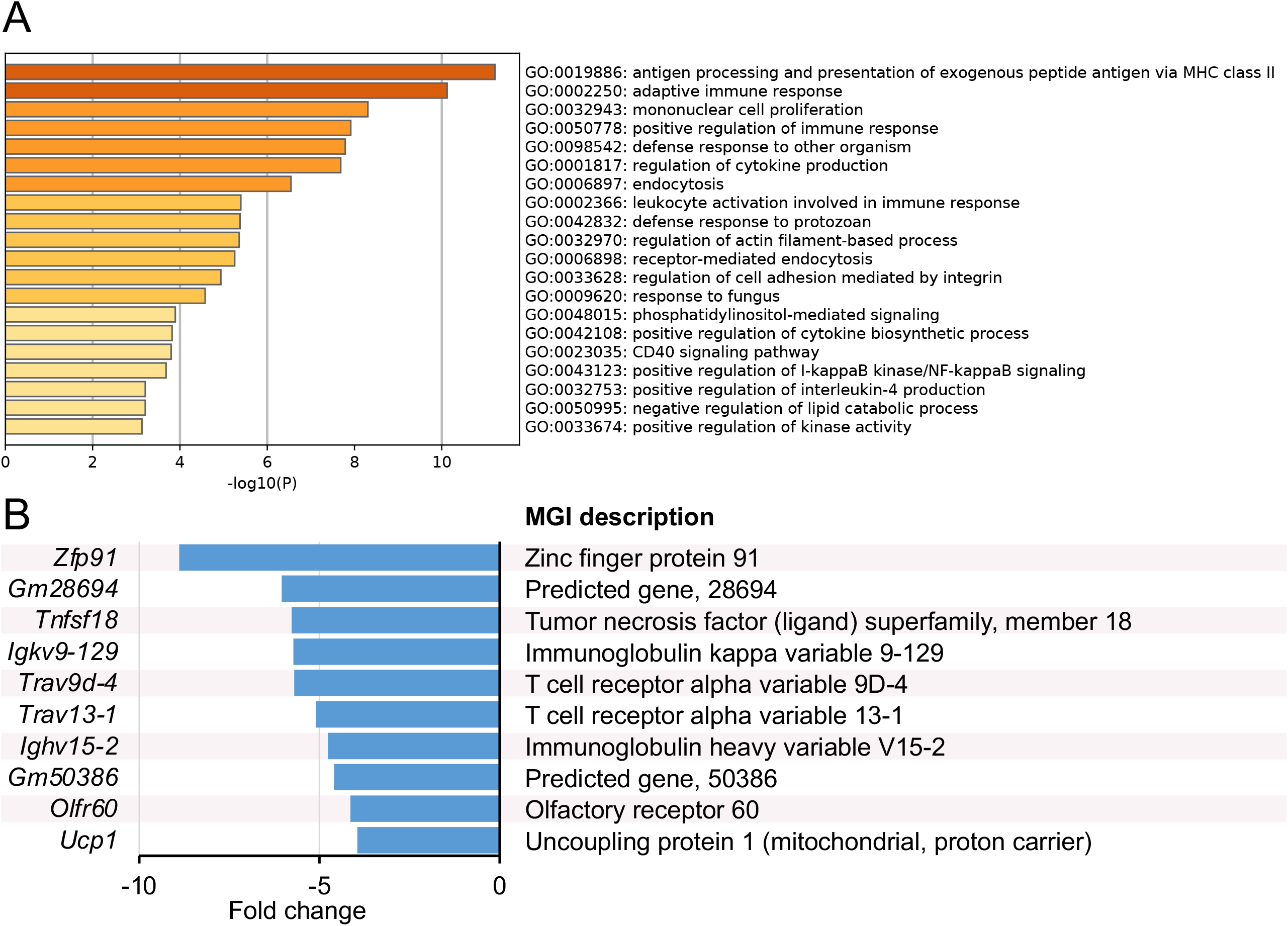
Gene ontology (GO) enrichment analysis of downregulated DEGs. (**A**) A heat map of GO terms across the upregulated DEGs list, colored to indicate the p values. Meta scape were used for this analysis. (**B**) Selected 10 genes indicating low false discovery rate of down regulated DEGs.

## Discussion

Hypothalamic neuropeptides regulate feeding behavior and systemic metabolism, are closely associated with obesity development^32,33^. We have recently shown that two-months overexpression of *Npgl*, a novel small secretory protein precursor gene, increased food intake and considerable fat accumulation in mice without metabolic abnormalities^27^. However, the underlying mechanisms of fat accumulation and maintaining metabolic normality in *Npgl*-overexpressing mice remain unknown. In this study, we performed *Npgl* overexpression for 18 days as the onset of obesity to address the mechanisms underlying fat accumulation and avoid secondary effects caused by prolonged gene overexpression. Our data showed that *Npgl* overexpression for 18 days was sufficient to induce fat accumulation with no metabolic abnormalities, such as hyperglycemia and hyperlipidemia. Furthermore, transcriptome analysis using RNA-seq revealed that *Npgl* overexpression upregulated the genes involved in cytoskeleton regulation, whereas it downregulated those involved in the immune-inflammatory response in the iWAT.

To date, considerable research has demonstrated that excess fat accumulation during obesity is accompanied by chronic inflammation and induces metabolic abnormalities, such as hyperglycemia, glucose intolerance, and insulin resistance^34,35^. On the other hand, in this study, transcriptome analysis indicated that *Npgl* overexpression could suppress the immune-inflammatory response in the adipose tissue, even though it induced considerable fat accumulation. It is known that immune cells in adipose tissues, such as adipose tissue macrophages (ATMs), contribute to the development of metabolic abnormalities^6^. For instance, continuous exposure to a high-fat diet induces fat accumulation and infiltration of pro-inflammatory M1 macrophages into adipose tissues^36,37^. M1 macrophages secrete pro-inflammatory cytokines, such as TNFα, resulting in inflammatory responses and subsequent metabolic abnormalities as well as lipolysis in adipocyte^36,38^. Our recent study has shown that *Npgl* overexpression for two months decreases mRNA expression of *Tnfα* and increases that of anti-inflammatory *adiponectin*, implying that NPGL can possess anti-inflammatory effects on adipose tissues^27^. Indeed, we have already demonstrated that *Npgl* overexpression improves glucose intolerance, insulin resistance, and hyperglycemia^39^. As well as ATMs, T cells in adipose tissue affect the inflammatory response during obesity development^4^. Studies in obese mice have shown that T cells accumulate in obese adipose tissue and short-term depletion of T cells improved systemic insulin resistance^40,41^. In the present study, we observed decreased mRNA expression of *Tnfsf18*, contributing to T cell activation. On the other hand, recent studies have revealed that there are a lot of small subpopulations of T cells in adipose tissues, regulating not only specific inflammatory response but also lipid metabolism during obesity development^42–44^. Further study to investigate the effects of NPGL on these immune cells will help clarify the mechanisms of fat accumulation and metabolic normality in *Npgl*-overexpressing mice.

In addition to the anti-inflammatory effects of NPGL, the present study demonstrated that *Npgl* overexpression upregulated genes involved in the cytoskeleton. We have demonstrated that *Npgl* overexpression enlarges adipocytes according to fat accumulation in rodents^24,27^. Several studies have shown that cytoskeleton remodeling is required to induced adipogenesis during fat accumulation^45,46^. Therefore, it is suggested that NPGL can play a role in cytoskeletal regulation to prepare subsequent fat accumulation. On the other hand, we observed increased blood insulin levels in *Npgl*-overexpressing mice. To date, several peripheral factors have been identified as incretins^47^. For instance, glucagon-like peptide-1, a 31-amino-acid hormone secreted from the lower intestine and colon, acts directly on the pancreatic islets to stimulate insulin secretion^48,49^. Moreover, secreted insulin promotes glucose uptake into peripheral tissue and subsequent adipogenesis^50^. Thus, NPGL may act as an incretin secreted from the hypothalamus and induce fat accumulation in the WAT via insulin signaling, even though *Npgl*-overexpressing for 18 days did not affect the activity of *de novo* lipogenesis.

In summary, this study revealed that *Npgl* overexpression for 18 days was sufficient to increase cumulative food intake and the masses of WAT without metabolic abnormalities, such as hyperglycemia and hyperlipidemia. Furthermore, transcriptome analysis of the iWAT using RNA-seq revealed that overexpression of *Npgl* upregulated the genes involved in cytoskeleton regulation, whereas it decreased those involved in immune-inflammatory responses. These results suggest that short-term *Npgl* overexpression plays a crucial role in enlarging adipocytes and suppressing inflammation to avoid metabolic abnormalities, eventually contributing to accelerating energy storage.

## Materials and Methods

### Animals

Male C57BL/6J mice (7 weeks old) were purchased from Charles River Laboratories (Kanagawa, Japan) and housed in standard conditions (25 ± 1°C under a 12-h light/dark cycle) with *ad libitum* access to water and normal chow (CE-2; CLEA Japan, Tokyo, Japan).

### Production of AAV-based vectors

AAV-based vectors were produced following a previously reported method^24^. In the present study, the primers for mouse *Npgl* were 5′ CGATCGATACCATGGCTGATCCTGGGC 3′ (sense) and 5′ CGGAATTCTTATTTTCTCTTTACTTCCAGC 3′ (antisense). The AAV-based vectors were prepared at a concentration of 1 × 10^9^ particles/µL and stored at −80°C until use.

### Stereotaxic surgery

For *Npgl* overexpression, mice were bilaterally injected with 0.5 µL/site (5.0 × 10^8^ particles/site) of AAV-based vectors that either carried the *Npgl* gene (AAV-NPGL) or served as controls (AAV-CTL). Vectors were injected into the MBH region (2.2 mm caudal to the bregma, 0.25 mm lateral to the midline, and 5.8 mm ventral to the skull surface) using a Neuros Syringe (7001 KH; Hamilton, Reno, NV, USA). Food intake and body mass were measured everyday (9:00 a.m.). Tissue and organ mass and blood biomarkers were assessed at the experimental endpoint.

### Quantitative RT-PCR

The iWAT and the liver were dissected from mice and snap-frozen in liquid nitrogen for RNA processing. Total RNA was extracted using TRIzol reagent (Life Technologies, Carlsbad, CA, USA; liver) or QIAzol lysis reagent (QIAGEN, Venlo, Netherlands; iWAT) in accordance with the manufacturers’ instructions. First-strand cDNA was synthesized from total RNA using a ReverTra Ace kit (TOYOBO, Osaka, Japan).

The primer sequences used in this study are listed in Table 1. The qRT-PCR was conducted following previously reported methods^24,26^. Relative expression of each gene was determined by the 2^−ΔΔCt^ method; the beta-actin gene (*Actb*) was used as an internal control for the liver, and the ribosomal protein S18 gene (*Rps18*) was used as an internal control for the iWAT.

**Table 1.**
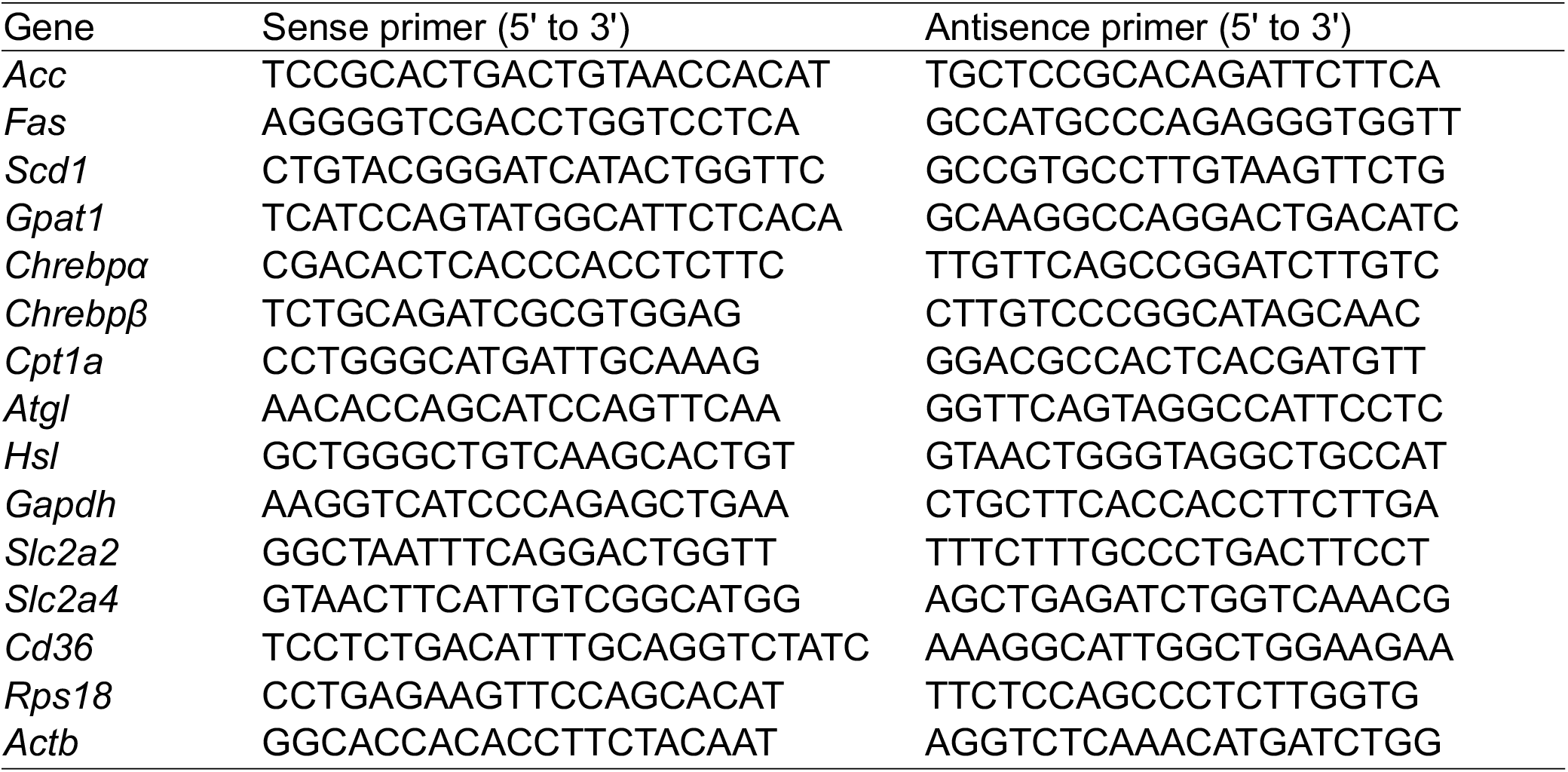
Primers for qRT-PCR used in this study

### Fatty acid analysis

For the analysis of endogenous SCD1 activity in the iWAT, the lipids were extracted according to the previous method^51^. Briefly, The iWAT (50 mg) was extracted with 500 µl of chloroform: methanol (2:1) using beads crusher (μT-12; TAITEC, Saitama, Japan) and 125 µl of distilled water was added and mixed by inversion. After incubation for 30 min, the sample was centrifuged at 3000 × g and the lower organic phase was collected and evaporated. Extracted fatty acids were methylated using Fatty Acid Methylation Kit (nacalai tesque, Kyoto, Japan) and purified using Fatty Acid Methyl Ester Purification Kit (nacalai tesque). The eluted solution was evaporated to dryness and kept at –20°C. The residues were resolved into hexane and fatty acids were identified by GC-MS (JMS-T100 GCV; JEOL, Tokyo, Japan). The SCD1 activity was estimated as oleate to stearate ratio (18:1/18:0) and palmitoleate to palmitate ratio (16:1/16:0) from individual fatty acids. The 16:1/16:0 ratio seems to be a better indicator of endogenous SCD1 activity than the 18:1/18:0 ratio^28^. The *de novo* lipogenesis index was calculated from palmitic to linoleic ratio (16:0/18:2n-6) from individual fatty acids^29,30^.

### Blood biomarker analysis

Serum levels of glucose, lipids, and hormones were measured using appropriate equipment, reagents, and kits. The GLUCOCARD G+ meter was used to measure glucose content (Arkray, Kyoto, Japan). The NEFA C-Test Wako (Wako Pure Chemical Industries, Osaka, Japan) was used to measure free fatty acid levels. The Triglyceride E-Test Wako (Wako Pure Chemical Industries) was used to measure triglyceride levels and the Cholesterol E-Test Wako (Wako Pure Chemical Industries) for cholesterol content. The Rebis Insulin-mouse T ELISA kit (Shibayagi, Gunma, Japan) was used to measure insulin levels. The Leptin ELISA Kit (Morinaga Institute of Biological Science, Yokohama, Japan) was used to measure leptin levels.

### Transcriptome analysis

Total RNA of the iWAT was extracted using QIAzol lysis reagent (QIAGEN). RNA-seq libraries were prepared and sequenced (150bp, paired-end) on the MGI DNBSEQ platform at GENEWIZ (Saitama, Japan). The sequence data (FASTQ files) were subjected to trimming process and evaluated by Trim Galore (https://www.bioinformatics.babraham.ac.uk/projects/trim_galore/). The mouse reference sequence file downloaded from HISAT2 ftp site (ftp://ftp.ccb.jhu.edu/pub/infphilo/hisat2/data), and the annotated general feature format (gff) file was downloaded from the GENCODE site (https://www.gencodegenes.org/mouse/release_M9.html). The RNA-seq reads were mapped to the reference genomic sequence by HISAT2 (https://ccb.jhu.edu/software/hisat2/manual.shtml) and then sorted by SAMtools (http://www.htslib.org/download/). StringTie (https://ccb.jhu.edu/software/stringtie/) processed the read alignments, estimates abundances where necessary, and creates new transcript tables. TCC-GUI compared all transcripts across conditions and produced DEGs^31^. Metascape (http://metascape.org/) was used for the GO enrichment analysis. A gene list for Metascape analysis was generated TCC-GUI, where genes were identified as ‘significantly differentially expressed’ (FDR < 0.1).

### Statistical analysis

Group differences between the mice injected with AAV-NPGL and those injected with AAV-CTL were statistically evaluated using Student’s t-test; *p* values < 0.05 were considered significant. Cumulative food intake and body mass were compared between the two groups at same day using the unpaired two-tailed Student’s *t*-test (Figure 1).

## Acknowledgements

We are grateful to Mr. Takaya Saito (Hiroshima University) and Mr. Atsuki Kadota (Hiroshima University) for the experimental support.

## Author Contributions

Conceptualization, K.F. and K.U.; methodology, K.F., Y.N., E.I-U., and M.F.; investigation, K.F., Y.N., E.I-U., M.F., H.B., and K.U.; writing—original draft preparation, K.F.; writing—review and editing, K.F., H.B., and K.U.; visualization, K.F.; project administration, K.U.; funding acquisition, K.F., E.I.-U., and K.U. All authors have read the manuscript and agreed to its published version.

## Statement of Ethics

All animal experiments were performed according to the Guide for the Care and Use of Laboratory Animals prepared by Hiroshima University (Higashi-Hiroshima, Japan), and these procedures were approved by the Institutional Animal Care and Use Committee of Hiroshima University (permit numbers: C13-12, C13-17, and C21-1-2).

## Informed Consent Statement

Not applicable.

## Data availability Statement

No big data repositories needed. The raw data supporting the findings of this manuscript will be made available by the corresponding authors, K.F., and K.U., to any qualified researchers upon reasonable request.

