## Supplementary Figure for "Neurosecretory protein GL-induced fat accumulation is accompanied by repressing the immune-inflammatory response in the adipose tissue of mice"

Supplemental Fig. 1

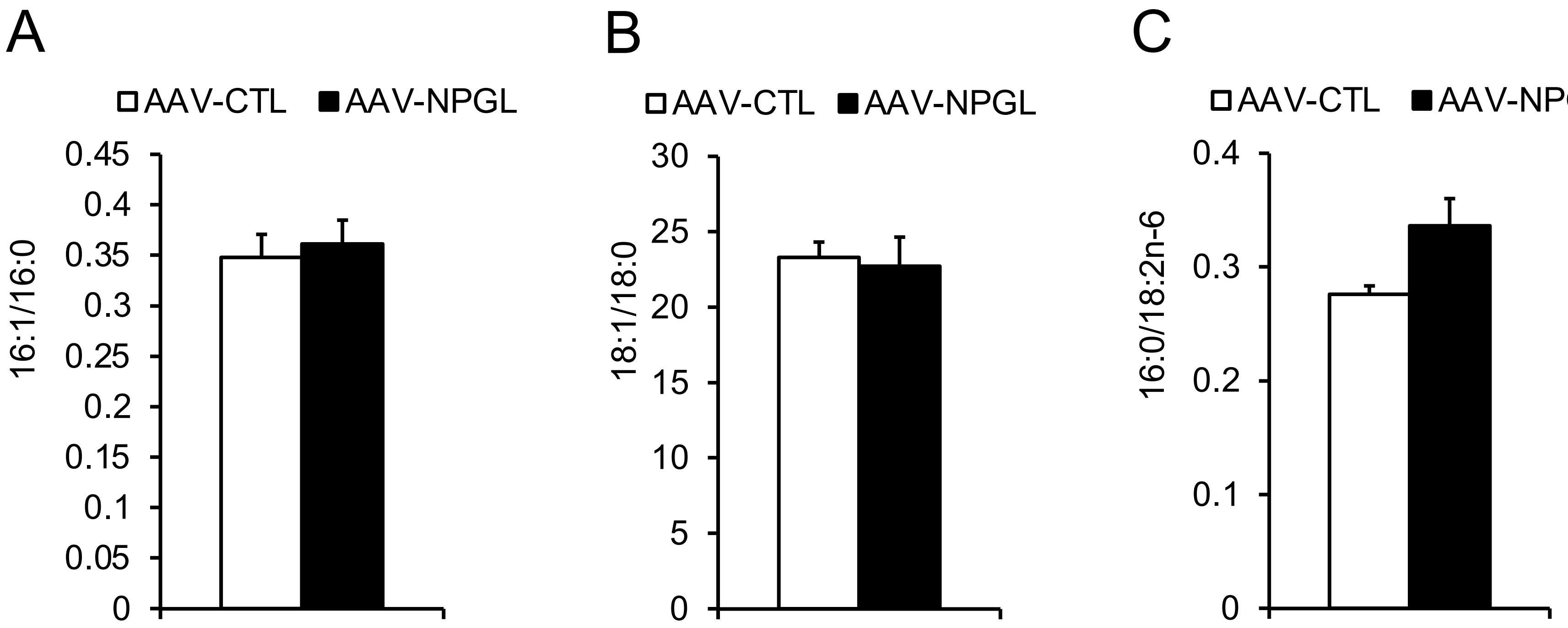


Supplemental Figure 1. Effects of Npgl overexpression for 18 days on fatty acid ratio in the in the inguinal white adipose tissue (iWAT). These mice were injected with an adeno-associated virus (AAV) vector, either a control (AAV-CTL) or a vector carrying the NPGL precursor gene (AAV-NPGL). (A) Fatty acids ratio (16:1/16:0). (B) Fatty acids ratio (18:1/18:0). (C) Fatty acids ratio (16:0/18:2n-6). Each value represents the mean ± standard error of the mean (n = 5–6).
